# Microclimate predicts frost-hardiness of alpine *Arabidopsis thaliana* populations better than altitude because the microclimate effect increases with altitude

**DOI:** 10.1101/544791

**Authors:** Christian Lampei, Jörg Wunder, Thomas Wilhalm, Karl J. Schmid

## Abstract

- In mountain regions average temperatures decrease at higher altitudes. In addition, microenvironmental conditions can strongly affect microclimate and may counteract average effects of altitude.
- We investigated winter frost hardiness of *Arabidopsis thaliana* accessions originating from 13 sites along altitudinal gradients in the Southern Alps during three winters on an experimental field station on the Swabian Jura and compared levels of frost damage with the observed number of frost days (<1°C) in eight collection sites.
- We found that frost-hardiness increased with altitude in a log-linear fashion. This is consistent with adaptation to higher frequency of frost conditions, but also indicates a decreasing rate of change in frost hardiness with increasing altitude. Moreover, the number of frost days measured with temperature loggers at the original collection sites correlated much better with frost-hardiness than the altitude of collection sites, suggesting that populations were adapted to their local microclimate. Notably, the variance in frost days across sites increased exponentially with altitude.
- Together, our results suggest that strong microclimate heterogeneity of high alpine environments may preserve functional genetic diversity in small populations. This challenges the suitability of habitat predictions based on large scale climatic variables (or proxies, such as altitude) for topographically complex areas.

## Introduction

In sessile organisms like plants local adaptation along environmental gradients such as aridity or temperature gradients may lead to clinal trait variation. For example, seed dormancy increases linearly with decreasing rainfall in many annual dryland species (Hacker, 1984; Hacker & Ratcliff, 1989; Volis *et al*., 2002; Kronholm *et al*., 2012; Wagmann *et al*., 2012; Tielbörger *et al*., 2012; Lampei *et al*., 2017). Mostly, environmental gradients result from geographic or geological conditions (e.g. altitude differences or rain shadow) that influence local climatic conditions. However, while geography may cause gradual changes in environmental variables at large spatial scale, microtopography may confound it at small spatial scale, resulting in a heterogeneous microclimate landscape. Exposure, snow cover, soil type and soil depth are just a few potential confounding factors which often vary on a local scale. For these reasons, geographic gradients such as latitudinal or altitudinal gradients are often poor proxies for the continuous change of single environmental factors (Körner, 2007; Graae *et al*., 2012; De Frenne *et al*., 2013). These effects of microtopography may severely complicate the modeling of climate change effects for the prediction of species distribution ranges (Dobrowski, 2011a; Graae *et al*., 2012; Oldfather & Ackerly, 2018). For example, migration to higher altitude may not be sufficient to follow the climatic niche because soil temperatures may be decoupled from air temperatures. On the other hand, a heterogenous microclimate landscape may allow species to follow their climatic niche by migrating just “around the corner” (i.e. change exposition) instead of moving at larger geographical scales. Thus, it is important to understand how microclimate changes along altitudinal gradients and how this affects the adaptation and distribution of plants.

One factor that changes with higher altitude is temperature. On average, atmospheric temperature drops by 5.5 K per 1000 m altitude (Körner, 2007). This results in a shift of growing season with vegetative growth starting later at higher altitudes, which partly offsets the average temperature difference for plants. Nevertheless, at high altitudes frequent and quick weather changes may lead to sudden frost periods. This explains why the ability to survive frost events is a typical adaptation to climate at higher altitudes (Sakai & Otsuka, 1970). For instance, in the central European Alps species with a higher upper distribution boundary showed increased summer frost resistance (Taschler & Neuner, 2004). Also, Andean forbs and grasses from a high-altitude site at 3600 m showed higher frost resistance than conspecifics from a lower site at 2800 m (Sierra-Almeida *et al*., 2009). However, higher altitude populations are not always more resistant to frost. In Sweden, *Pinus sylvestris* showed higher frost resistance with higher latitude, but not with higher altitude (Sundblad & Andersson, 1995). We previously showed that Southern Alpine *A. thaliana* populations from 2,200-2,350 m were not better adapted to frost experience than valley populations from 600-1,000 m altitude despite significant variation in frost-hardiness among populations (Günther *et al*., 2016). In this study we argued that microclimatic effects together with a delayed start of the growing season may reduce the risk of frost damage at higher altitude sites. According to the “law of the relative constancy of habitat” a species would shift its habitat niche towards warmer micro-sites when the climate is cold in relation to its core habitat (Walter & Walter, 1953). This raises the question to which extent climatic conditions at higher altitude select for higher freezing tolerance and to which extent plants use sites with a warm microclimate, or “microrefugia” (Dobrowski, 2011b), to avoid freezing damage.

To study this question, we tested frost-hardiness of *A. thaliana* plants from Southern Alps in the provinces Bolzano and Trento, collected from sites ranging in altitude from 280 to 2,355 meters above sea level. Plants were subjected to winter freezing under near natural conditions during three winters on the Swabian Jura. Additionally, we monitored top soil temperatures over two years at eight sites with data loggers. Combining these data sets, we compared the effect of altitude with the pure effect of freezing probability on the evolved frost-hardiness of plants. As annual species *A. thaliana* is part of an underrepresented category in the high alpine flora. Nevertheless, *A. thaliana* was found up to an altitude of 4,200 m a.s.l. (Al-Shehbaz & O’Kane, 2002; Zeng *et al*., 2017). There is some evidence that *A. thaliana* populations can adapt to high altitudes. In a genome scan, Kubota et al. (2015) found convergent differentiation of genomic regions that were associated with ecological relevant parameters in *A. thaliana* plants from altitude transects on two independent mountains. Also, in a common garden study, high-altitude populations of the Eastern Pyrenees showed higher aboveground biomass and increased fecundity, suggesting selection for higher vigor (Montesinos-Navarro *et al*., 2011). In contrast, accessions from high-altitude populations in Switzerland were smaller and showed a reduced vigor across three common gardens at different altitudes (Luo *et al*., 2015), although a dwarf accession showed increased fitness at high altitudes. Vidigal et al. (2016) found that seed dormancy decreased, seed size increased and plants flowered later with higher altitude of the collection site on the Iberian Peninsula. In populations from the North Italian Alps, a high differentiation of genomic regions with annotations related to ecological relevant parameters such as soil conditions, pathogen response or soil and light response was observed (Günther *et al*., 2016). However, geographic patterns of the traits frost resistance, UV-B and light stress response did not suggest adaptation to high altitude (Günther *et al*., 2016). So far, we are not aware of any other study that studied frost-hardiness of *A. thaliana* population from different altitudes. Tests with low altitude accessions of *A. thaliana* suggest that the species avoids freezing via super cooling (Reyes-Díaz *et al*., 2006), although freezing tolerance also plays an important role. Zhen and Ungerer (2008) showed that accessions varied in their freezing tolerance in a clinal fashion with increased freezing tolerance at high latitudes. In our study we tested the following predictions: (1) Frost-hardiness of *A. thaliana* increases with altitude. (2) The average frequency of frost explains frost-hardiness better than altitude. (3) Altitude is a better predictor of frost frequency at low-altitude sites than at high altitudes. To test these hypotheses, we included a larger number of populations from the Southern Alps collected at different altitudes and microsites to compare the role of different spatial scales in adaptation to altitude.

## Materials and Methods

### Plant material

We collected and geotagged seeds of *Arabidopsis thaliana* accessions in the Southern Alpine provinces of Bolzano and Trento during summers from 2006 to 2008 (Fig. 1, Table 1). Also, we collected accessions from new micro-sites within the Finail site in summer 2011 to increase the sample size from this location. Local populations at the collection sites differed considerably in size, which is mirrored in our experiments (Table 1). We reared and self-fertilized plants under standard greenhouse conditions to produce S1 seeds for the experiments.

**Fig. 1.**
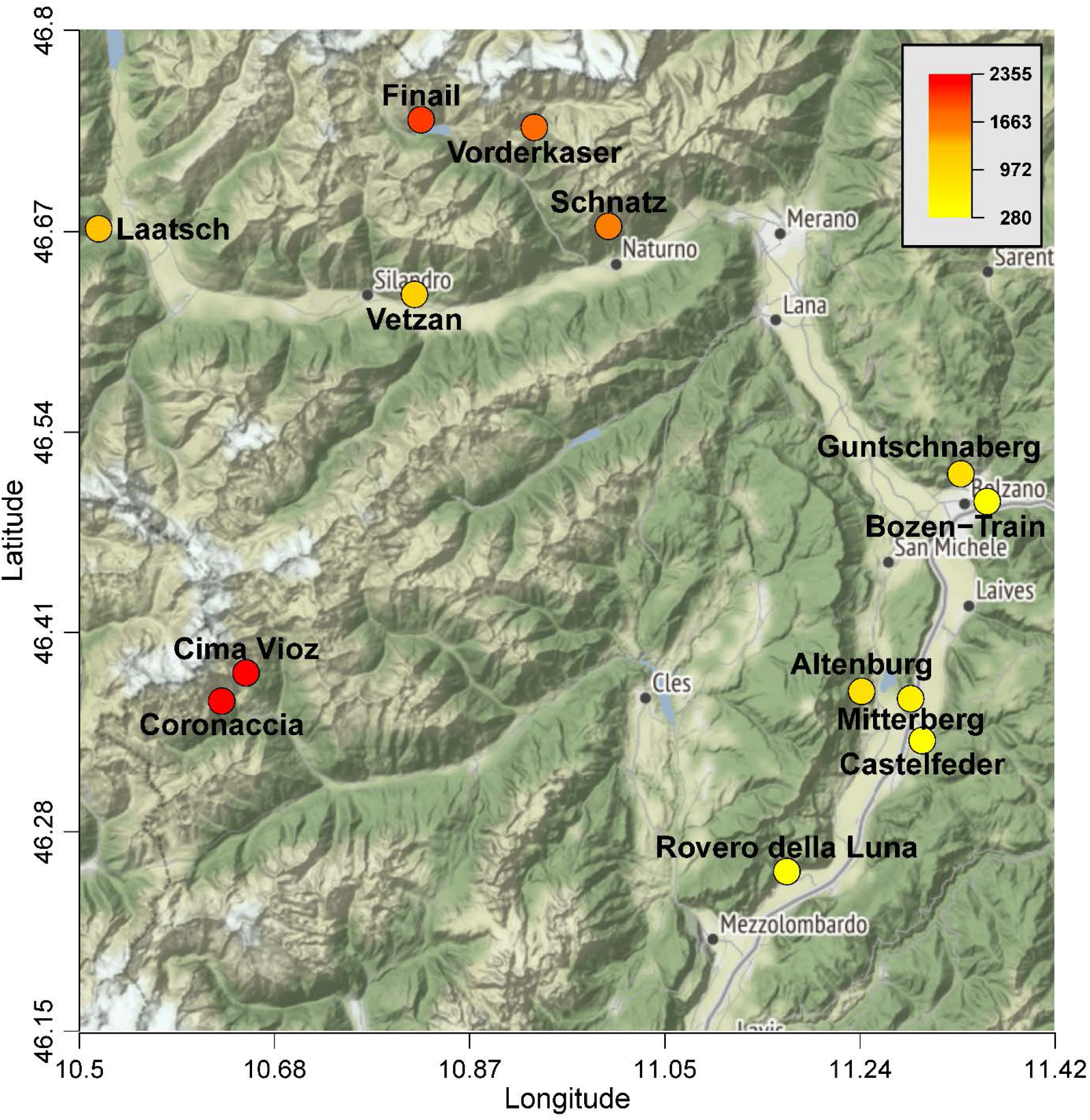
Map of the *A. thaliana* collection sites in the Southern Alpine provinces of Bolzano and Trento with heat color indicating the altitude of each site. Map source: http://maps.stamen.com/terrain/?request=GetCapabilities&service=WMTS&version=1.0.0#11/46.5223/10.9980.

**Table 1.** Characterization of sites, numbers of *A. thaliana* accessions tested and period of logged temperatures. In brackets province codes (BZ = Bolzano, TN = Trento).

| Site | Altitude<br>(m) | Exposition | Disturbance<br>(human) | Grazing/<br>trampling | Accessions<br>2010/2012 | Collection<br>year | Temperature<br>Logged |
| --- | --- | --- | --- | --- | --- | --- | --- |
| Altenburg (BZ) | 625 | Flat | strong | non | 8/4 | 2008 | - |
| Guntschnaberg (BZ) | 730 | SE | little | non | 8/4 | 2006 | - |
| Bozen-Train (BZ) | 280 | Flat | strong | non | 3/3 | 2006 | - |
| Castelfeder (BZ) | 390 | SE-SW | intermediate | non | 12/4 | 2008 | 2012-15 |
| Cima Vioz (TN) | 2355 | S | undisturbed | chamois | 1/4 | 2007, 2012 | 2012-13 |
| Coronaccia (TN) | 2210 | SSW | undisturbed | chamois | 4/4 | 2007 | 2013-15 |
| Finail (BZ) | 2166 | SE | undisturbed | chamois | 6/4 | 2007, 2011 | 2012-15 |
| Laatsch (BZ) | 990 | SE | undisturbed | non | -/4 | 2007 | 2012-15 |
| Mitterberg (BZ) | 565 | S-SE | undisturbed | non | 8/4 | 2008 | 2012-15 |
| Roverè della Luna (TN) | 315 | Flat | intermediate | non | 1/1 | 2008 | - |
| Schnatz (BZ) | 1709 | SE | intermediate | Sheep | 1/1 | 2008 | 2012-15 |
| Vetzan (BZ) | 880 | S | little | non | 10/4 | 2008 | 2012-15 |
| Vorderkaser (BZ) | 1770 | SE | intermediate | cows | 3/3 | 2007 | 2012-13 |

### Experiments

In August 2010 we sowed seeds onto frost-hardiness testing tables situated on the experimental station “Oberer Lindenhof” of University Hohenheim on the Swabian Jura (48°28’24 N, 9°18’18 E, 720 m a.s.l.). The tables were 22 cm deep, 70 cm wide and 2.5 m long and their bottom was about 60 cm above the ground (Fig. 2a,b). We filled the tables with compost (pH=6.9, P=31.0 mg/100g, K=48.0 mg/100g, MgCaCh=5.6 mg/100g, humus=5.18 %) till about 5 cm below the top.

**Fig. 2.**
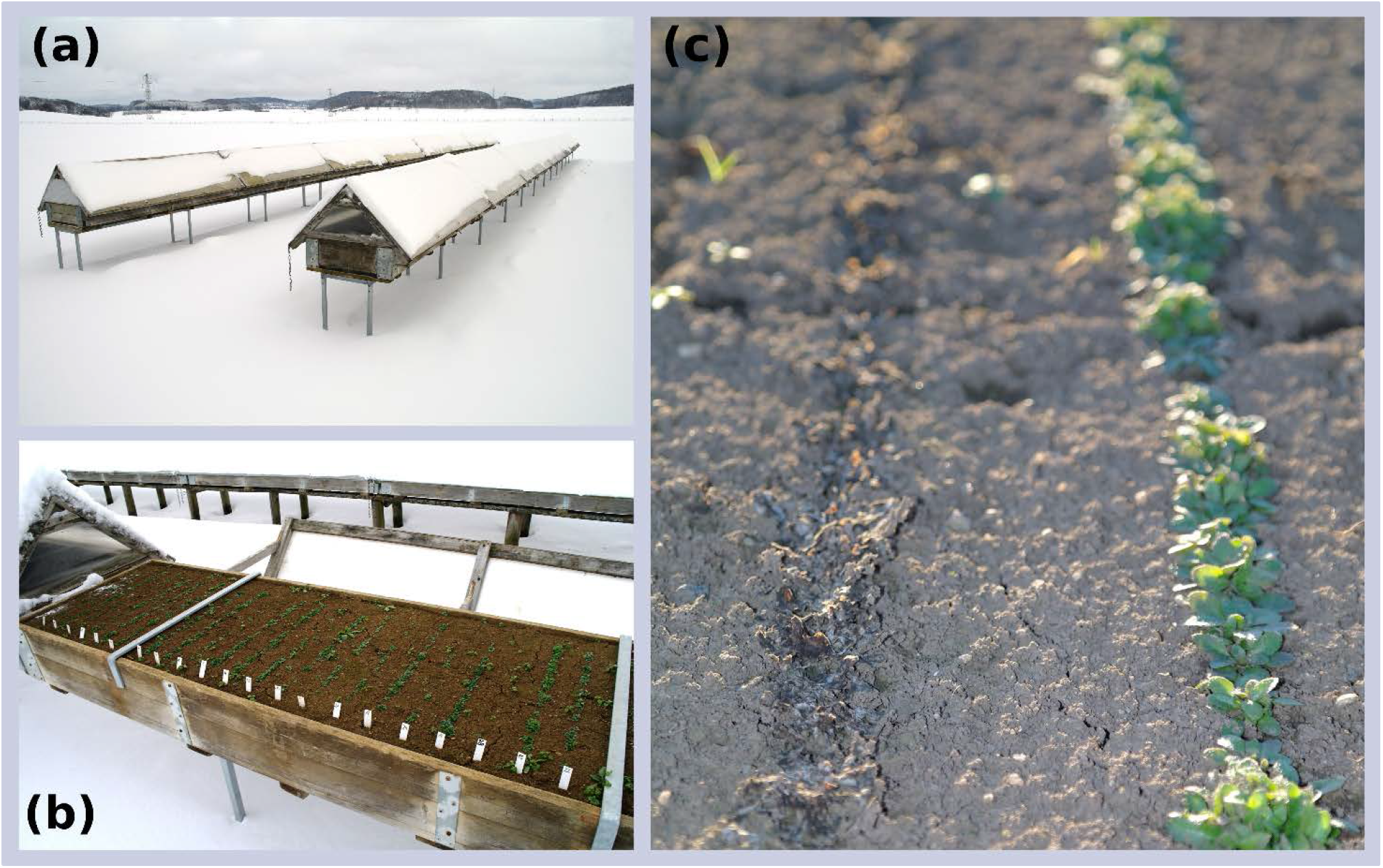
Frost testing boxes on the Swabian Jura and *A. thaliana* plants at the time of frost damage evaluation. (a) Transparent covers keep the snow from plants and poles ensure exposure to frost from all sides. (b) The plants appear unharmed when temperatures are below or close to zero. (c) The damages are visible after thawing of the damaged tissue. Here the left row was completely damaged while the right row survived almost unharmed.

Then we watered the soil extensively before adding another layer of moist humus that filled the tables to the top. Rows of seeds were parallel to the short edge of each table, 15 cm apart. We sowed each accession to two random rows (Fig. 2c), and number labeled them for blinded evaluation. Seeds were watered regularly to induce germination. After germination, we left plants to natural rainfall and irrigated only during dry periods to prevent plant loss by drought. On days with predicted snowfall during winter, a transparent cover that allowed air flow was put over the plants to keep them free of snow for direct frost-exposure (Fig. 2a, b).

In August 2012, we installed a second experiment on the frost-hardiness testing tables. This experiment was similar to the first one, but this time we used a more balanced experimental design, because in the first year, the number of accessions differed strongly among populations (Table 1). For a more even representation, we sowed four random accessions from each site of origin, if available, each with two replicates. If the number of available accessions was below four, the available accessions were replicated more often to reach eight replicates for each site.

In August 2013 we set up a third experiment on the frost-hardiness testing tables. This time, space was limited. Therefore, we used population replicates instead of individual accessions. We mixed equal amounts of seeds of four accessions for each site before sowing. Only in the sites Vorderkaser (*n*=3) and Schnatz (*n*=1) we had fewer accessions available. In this design, each of the six replicate rows resembled a true replicate for the collection site.

For each of the three experiments the daily minimum and maximum temperatures were obtained by the weather station of the experimental site (Fig. S1).

### Assessing frost damage

After the exposure of plants to natural winter frosts on the Swabian Jura, we evaluated the percent frost damaged tissue visually, while the accession identity was hidden from the monitoring person. Because frost damage is not visible while plants are still frozen, we let the plants recover and monitored frost damage some weeks after a period with strong frost (Fig. S1). At this time the affected tissue showed clear signs of decay (Fig. 2c). This time lag between frost damage and its evaluation is unlikely to bias our estimates due to the low growth rates during this time. A similar approach for assessing frost damage was used by Zhen and Ungerer (2008).

### Logging on-site soil temperature

In each site, we logged temperature using waterproof Hobo^®^ 8K Pendant^®^ Temperature Loggers from Onset^®^ (470 MacArthur Blvd., Bourne, MA 02532, USA). We submerged two temperature loggers in each site 3 cm in the ground to record the topsoil temperature with a 2 h resolution for the whole year. This way we exposed the data loggers to the same top-soil temperature that the plants experienced. Top-soil temperature is probably the most relevant temperature for plant survival because winter-annual *Arabidopsis thaliana* plants remain ground-bound rosettes until they start to reproduce in spring (Ågren & Schemske, 2012). We attached the data loggers to a nylon cord of 50-100 cm length which we covered by soil and tied to a nail that was fixed in the soil nearby a color marked shrub or rock, to facilitate recovery in the next summer. We recorded temperatures in nine sites that represent the altitudinal range covered by this study. For most sites we measured temperatures over a period of 2.5 years from July 2012 until January 2015, except for Cima Vioz and Vorderkaser (winter 2012/2013 only) and Coronaccia (July 2013 until January 2015). We also recorded micro-habitat specific temperatures in the two highest sites, Cima Vioz and Coronaccia. Here, *A. thaliana* plants grew underneath steep rock walls and in cracks in these rocks. We installed data loggers both below the rocks and in rock cracks that were large enough to support soil accumulation to ensure that soil temperature was measured.

### Analysis of temperature records

We trimmed temperature data from each data logger to cover two years starting and ending in mid-summer. For each data logger, we counted the number of days with temperatures below 1 °C, because our sample included sites in the valley where soil temperatures did not decrease below zero during winter. We averaged over all data loggers and recorded winters for each site to obtain an estimate for frost exposure of winter-annual plants. This estimate could be biased if the two winters were very different because we did not have records from both winters for all sites. Using the sites with full records, we fitted a generalized linear model with Poisson error family (corrected for overdispersion), to test if years differed across sites in their counts for days with temperatures below 1 °C (frost days). The model contained the dependent variable frost days and the independent variables year, site and their interaction. As expected, the number of frost days differed strongly between sites (F=125, df=5/14, p<0.001). However, there was no significant difference among years (F=1.29, df=1/19, p=0.28) and no interaction between sites and years (F=0.7, df=5/9, p=0.64). We concluded that the two winters were similar enough and required no correction.

### Frost damage

We evaluated frost damage as an average over all plants in a row (Fig. 2b, c). The estimated frost damage is then a measure of central tendency. This explains why binomial models fitted to the odd ratio of damaged over undamaged tissue showed very large overdispersion (dispersion parameter = 35.5) and the distribution of the residuals deviated strongly from normality. The central limit theorem states that the distribution of means quickly approaches normality with increasing sample size of each mean, independently of the distribution of the data. We therefore used percentage frost damage as dependent variable with a Gaussian error distribution instead, which considerably improved the fit of residuals with a normal distribution. To test if frost damage correlates with the altitude of collection sites, we fitted a linear mixed effects model with the following equation:

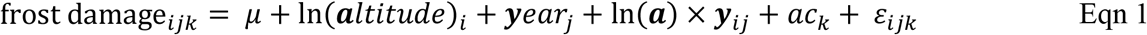

where μ is the overall intercept, ln(*altitude*)_*i*_ the fixed effect of the natural logarithm of altitude of population *i, year_j_* the fixed effect of year *j* and ln(*a*) ×*y_ij_* their interaction. With the random effect *ac_k_* of accession *k* we accounted for the non-independence of replicates of the same accession. The term *ε_ijk_* is the residual associated with the replicate of accession *k* from altitude *i* in the year *j*. We computed the model using the R package lme4 (version: 1.1-17; Bates et al. 2015). To test whether altitude or days below 1° C explained more variance in frost damage, we included this variable in the fixed effects part according to the following equation:

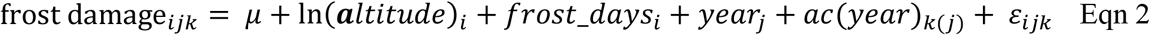

with the overall intercept *μ*, the logarithmic altitude effect of population *i*, the effect of the number of frost days of population *i*, and the random effects of year *j* and of accession *k* nested in year *j*.

The term *ε_ijk_* is the residual defined like in Eqn 1. The random effects were tested for models fitted with restricted maximum likelihood (REML) using either likelihood ratio tests by comparing two nested models or using the function ranova from the R package lmerTest (version: 3.01; Kuznetsova et al. 2017). For the fixed effects we refitted the models using maximum likelihood and applied Bayesian inference with a flat prior using the nsim-function in the R package arm (version: 1.10-1; Gelman and Yu-Sung 2018). As a measure of statistical significance, we present the Bayesian 95% credible interval (CrI 95%) for each estimate.

Further, we tested for a linear relationship between soil temperature (frost days) and altitude of the collection site using the lme function from the r-package nlme package in R (version: 3.1-137; Pinheiro et al. 2016), because it includes functions to account for heterogeneous variances. We fitted a mixed effects model of the form:

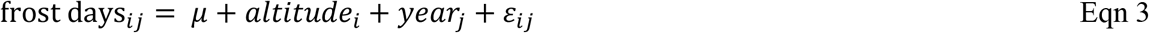

The model contained the overall intercept *μ*, the fixed effect of the altitude of site *i* and the random effect of year *j*. The term *ε_ij_* is the residual of the replicate in altitude *i* and year *j*. To model the variance proportionally to altitude we used the varPower function with the weights argument in the lme function. This means that the residuals had the form:

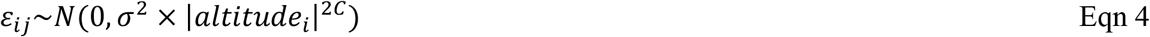

Here *N* is a normal distribution with mean zero and variance σ^2^ multiplied with the power of the absolute value of the altitude of site *i*. The variance function coefficient *C* is estimated from the data (Zuur *et al*., 2009).

## Results

### Accessions from higher altitude are more frost resistant

Our common garden experiment showed that frost damage significantly decreased with increasing altitude of the collection site if values were averaged across all three experiments (*b*=-19.2, CrI 95%: −32.8; −5.8). The three years differed strongly in frost damage, which was mostly due to strong variation in frost damage of the low altitude populations (Table 2). The variance of predicted frost damage across experiments at 280 m (σ^2^=827) was ten times the variance of frost damage estimates from 2,355 m altitude (σ^2^=80.4). Frost damage was strongest in the first winter (2010/2011; Fig. 3a), lower in the second winter (2012/13; Fig. 3b) and lowest in the third winter (2013/14; Fig. 3c). These differences between experiments match the number of relevant deep frost events (below −7 °C), in each winter. In a previous freezing experiment we found that the exothermic peak of freezing plant material of South Tyrolian *A. thaliana* plants was around −7 °C (Günther *et al*., 2016). The number of days with temperatures below −7 °C in the respective winters before frost damage was 18 in the first, 14 in the second and 2 in the last winter (Fig. S1). Consistent with the general frost damage the regression slope changed strongly between experiments (Table 2). Frost damage reduced dramatically with altitude of the collection site in the first winter, the slope was less steep in the second winter and insignificant in the last winter (Table 2). Notably, the model had been significantly improved by fitting frost damage to the natural logarithm of altitude, instead of assuming a simple linear relationship (χ^2^=4.8, df=0, p<0.001). This indicates that frost-hardiness showed a stronger correlation with altitude at the low end of the gradient than at its high end. To demonstrate the effect size of this change we calculated the frost damage increment for the lowest and the highest 500 m altitude segment (Table 2). The frost hardiness increment was 4.3-fold lower for high altitude populations than for populations in the lowest altitude segment. This indicates that the altitude difference between populations is less important for variation in frost hardiness at high altitude.

**Fig. 3.**
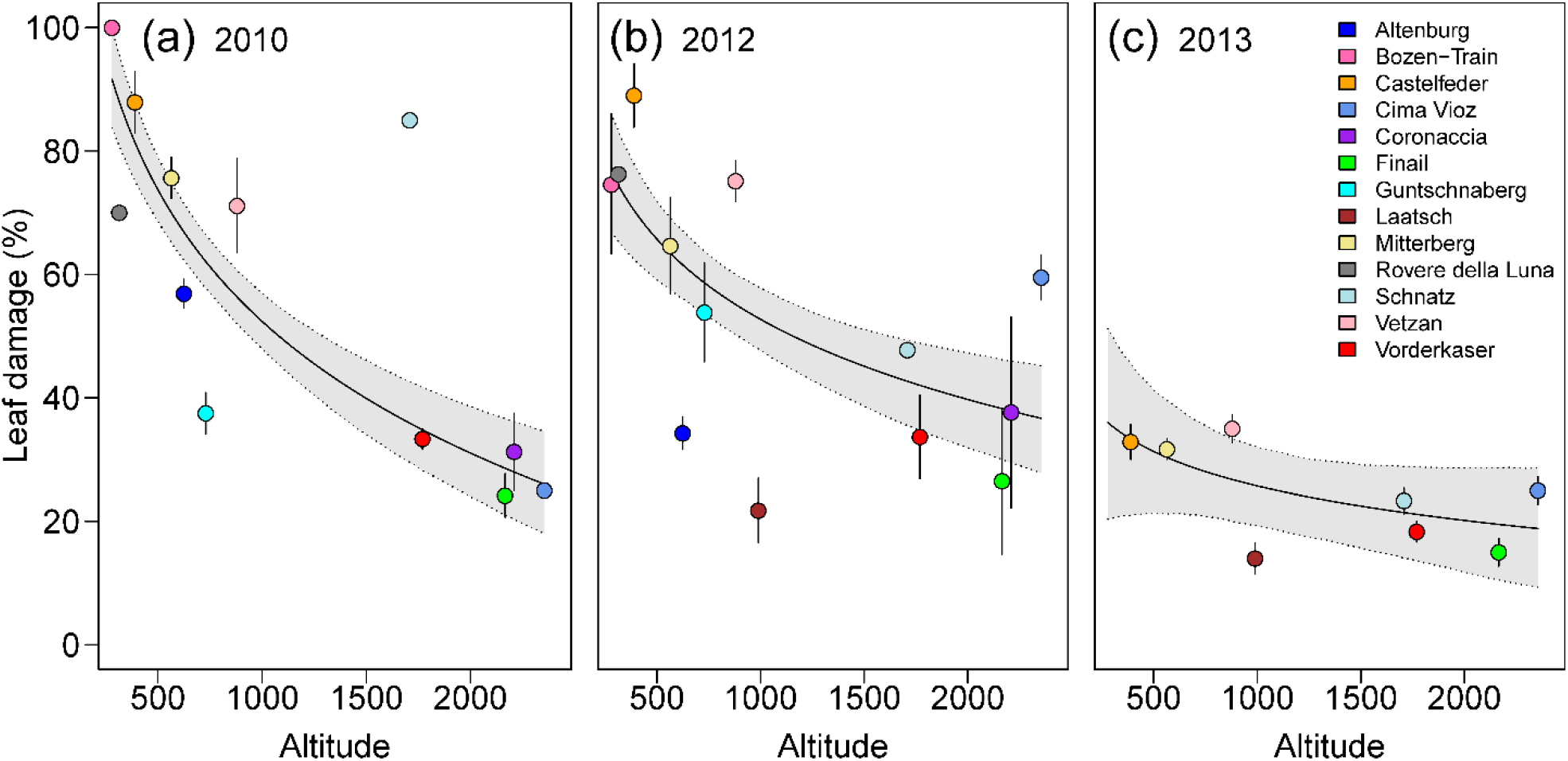
Frost damage of *A. thaliana* populations as a function of the altitude of the collection site. Populations from higher altitudes show reduced frost damage especially in (a) winter 2010/11, (b) winter 2012/13 and less in (c) winter 2013/14. The regression line is shown together with its 95% credible interval. In addition, for each site the averages and standard errors across accession means are presented. In the last year (c) the standard error was estimates across samples.

**Table 2.** Model predictions for the percentage of frost damaged leaf area and the change in frost damage with altitude for the high and the low altitude end of the gradient in the 3 experiments. The corresponding 95% credible intervals are presented in brackets.

| Experiment<br>(Winter) | Predicted frost damage |  | Predicted change in frost<br>damage | Predicted frost damage<br>increment |  |
| --- | --- | --- | --- | --- | --- |
|  | Low altitude<br>(280 m a.s.l.) | High altitude<br>(2,355 m a.s.l.) | Slope *b* | Lowest 500 m | Highest 500 m |
| 2010/11 | 92 (84; 99) | 26 (18; 34) | -31 (-37; -25) | -32 (-35; -28) | -7 (-9; -6) |
| 2012/13 | 77 (67; 86) | 37 (28; 45) | -19 (-34; -4) | -19 (-24; -14) | -4 (-6; -3) |
| 2013/14 | 36 (21; 51) | 18 (9; 29) | - 8 (-26; 11) | -8 (-16, 0.33) | -2 (-0.3; -4) |

### Top-soil temperature is a better predictor than altitude

To compare the effect of altitude with *in-situ* measured temperature, we reduced the dataset to the populations where the temperature was recorded and included the year as a random effect component (see methods for details). We found that the number of recorded frost days in the collection site (days with minimum temperature below 1° C) was a stronger predictor for differences in frost damage than the altitude of the collection site as can be seen from their partial regression slopes (Fig. 4). Also, when comparing two models that differed only in their fixed effects, the model with frost days was significantly better than the model with altitude (χ^2^=23.6, df=0, p<0.001), and in a model with both variables, frost days explained more variance (F=32.1) than altitude (F=4.7). Averaged across the three experiments the partial regression slope suggested a reduction in leaf damage of −0.28 (CrI 95%: −0.37; −0.18) with each additional frost day in the collection site of a population. In other words, if the number of frost days increased by 10 days from one to the next collection site, the frost damage of the respective plants in our common garden experiments decreased on average by 3%. The slope for altitude was −7 and was also significant (CrI 95%: −15; −0.66). These two slopes are not directly comparable, because altitude was log-transformed, showing that the change in frost damage was strong between valley populations and decreased toward high altitudes, as was found for the individual years (Table 2). As an additional test whether altitude or the number of frost days better explained the variation in frost damage we use the posterior distribution of 2,000 simulations to ask how likely it is to observe a slope equal to or greater than zero. While for altitude non-negative slopes were observed in 2% of the simulations, for frost days all slopes were negative, again demonstrating the superiority of frost days in explaining the observed differences between populations in frost damage.

**Fig. 4.**
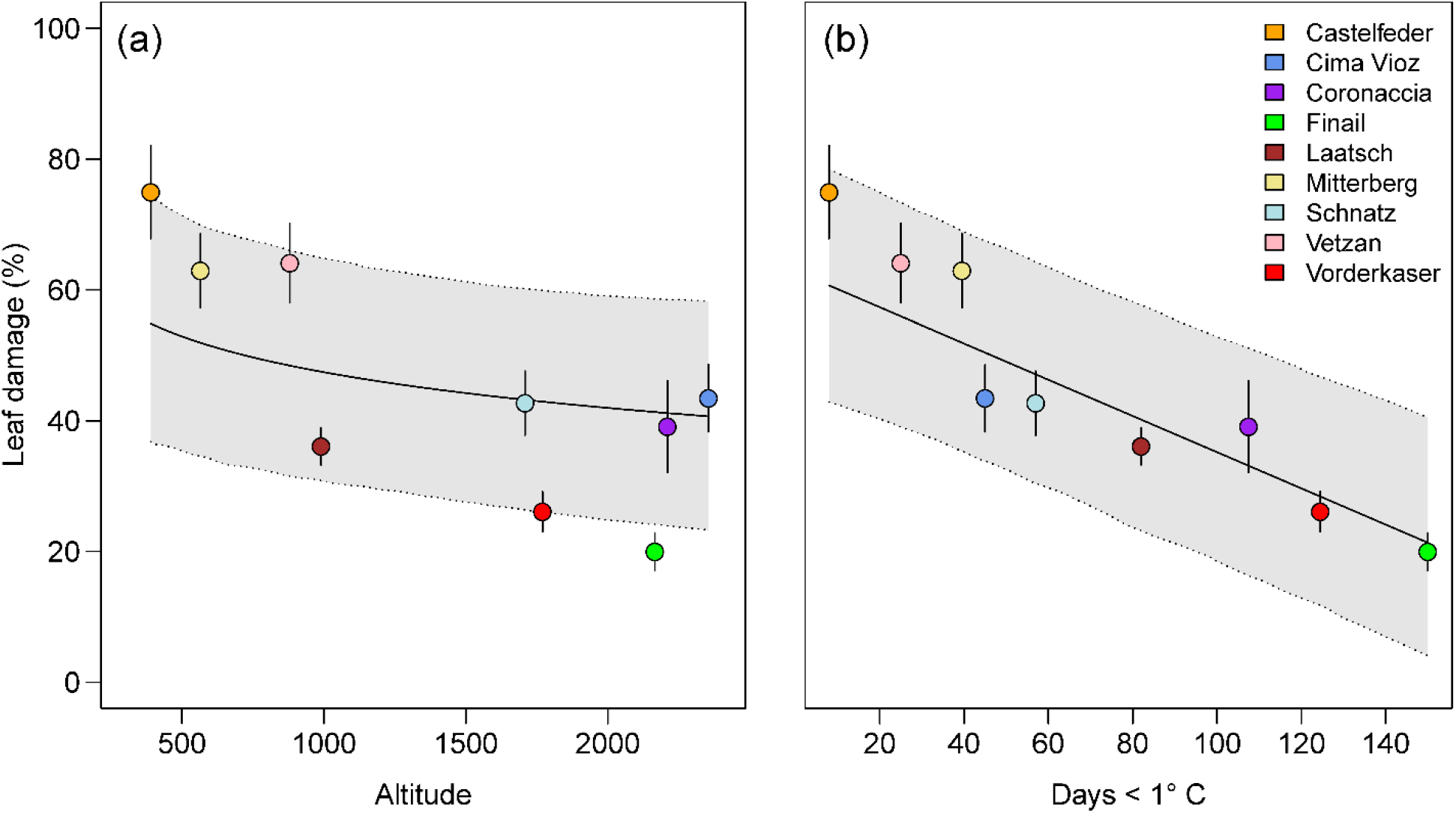
Partial regression lines and their 95% credible interval (grey) for frost damage of *A. thaliana* populations regressed on altitude (a) and frost days (b) averaged across the three experiments. Points represent population averages with standard error.

### Mismatch between altitude and soil temperature increases with altitude

The observation that the number of frost days explained variation in frost damage better than altitude, suggests that the probability to experience ground frost is not a simple function of altitude. To better understand how altitude influences the number of frost days we next tested whether there is a linear relationship between soil temperature and altitude of collection sites. Altitude indeed had a strong positive effect on the number of days below zero (Fig. 5a; F_1/24_=25.46, p<0.001), however the residual distribution was heterogeneous, as indicated by a significant Fligner-Killen test of homogeneity of variances (χ^2^=17.03, df=8, p=0.029). Notably, the variance strongly increased together with altitude as can be seen from the population averages displayed in Fig. 5a. We modeled this effect with a power function for the variance using the weights argument in the lme function (Pinheiro *et al*., 2016). As a result, the effect of altitude increased (F_1/24_=40.17, p<0.001) and the corrected residuals were distributed evenly. A likelihood ratio test showed that the correction of variance heterogeneity improved the model (ΔAICc=-7.5, L=10.5, df=1, p=0.001). The variance power function is plotted in Fig. 5b and suggests an exponential increase of residuals with altitude with a 15-fold increase in variance from 280 to 2,355 m altitude.

**Fig. 5.**
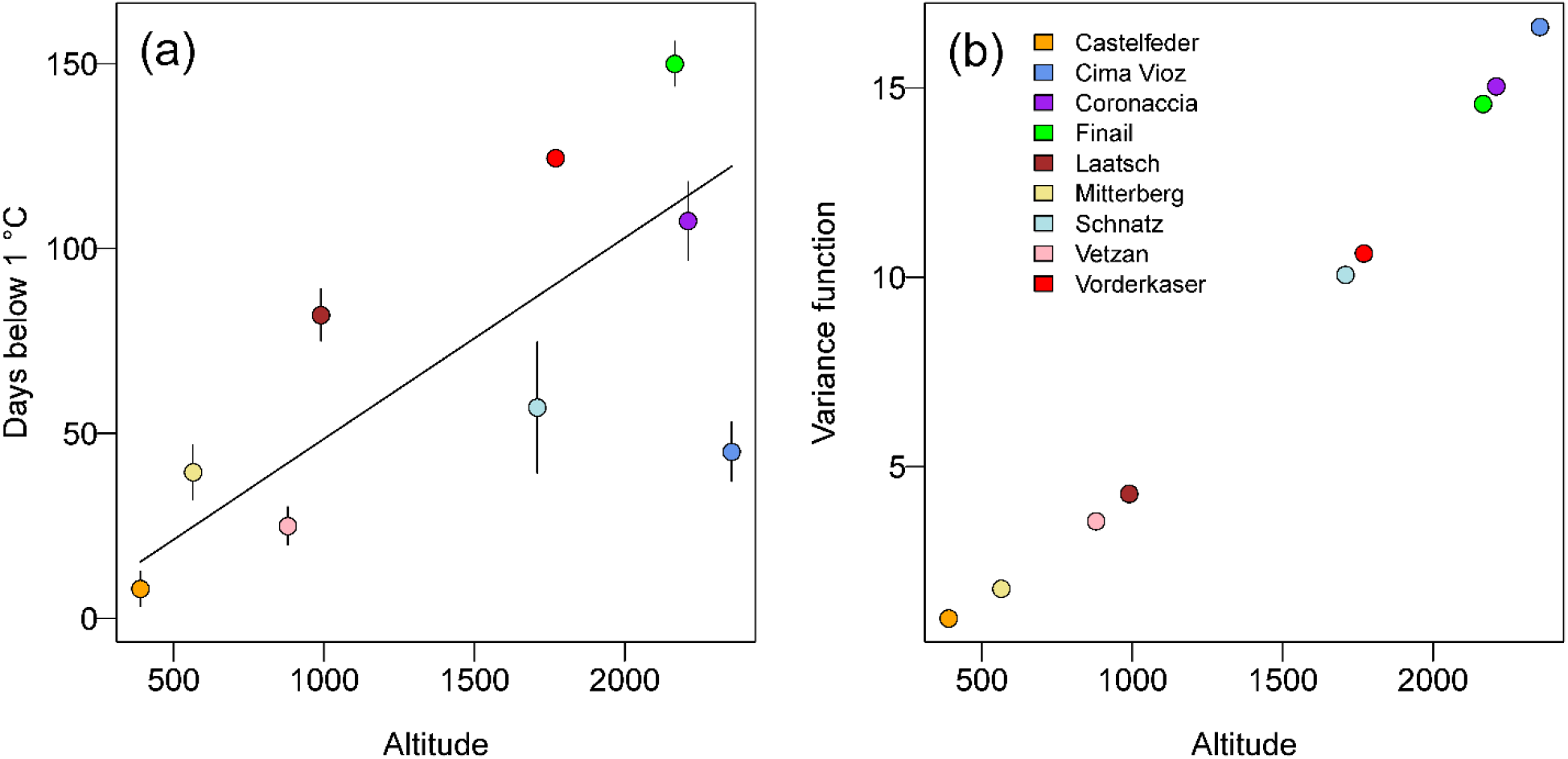
The effect of altitude on top-soil temperature. (a) Regression of frost days on altitude with site averages across data loggers and winters and standard errors. (b) Fitted values for the variance power function of the form *f_(altitude)_* = |*altitude_i_*|^(2x*C*)^ (*C* = variance function coefficient) that was used to model the increase of residuals with altitude.

To characterize the difference between two closely located microhabitats that harbored *A. thaliana* populations at the highest altitude we compared top-soil temperatures in rock cracks and below rocks (Fig. 6). In both sites the two microhabitats differed strongly in number of days with ground frost (i.e., days with temperature < 0°C). Here we counted the days with soil temperatures below zero because all temperature records included sufficient observations of ground frost. In Cima Vioz, we measured 29 ground frost days in the rock crack and 13 in the soil below the rocks during the winter 2012/2013. The Coronaccia site experienced 55 frost days in the rock crack and 82 in the soil below the rocks in the winter 2013/2014. Although measurements were taken in different winters, the daily maximum temperatures showed much higher variance on the ground (Cima Vioz: σ^2^=53.7; Coro: σ^2^=50.1) than in the rock crack (Cima Vioz: σ^2^=17.4; Coro: σ^2^=31.1) at both sites. Taken together, these data demonstrate that microsites indeed strongly differ in their temperature regime in the high mountains.

**Fig. 6.**
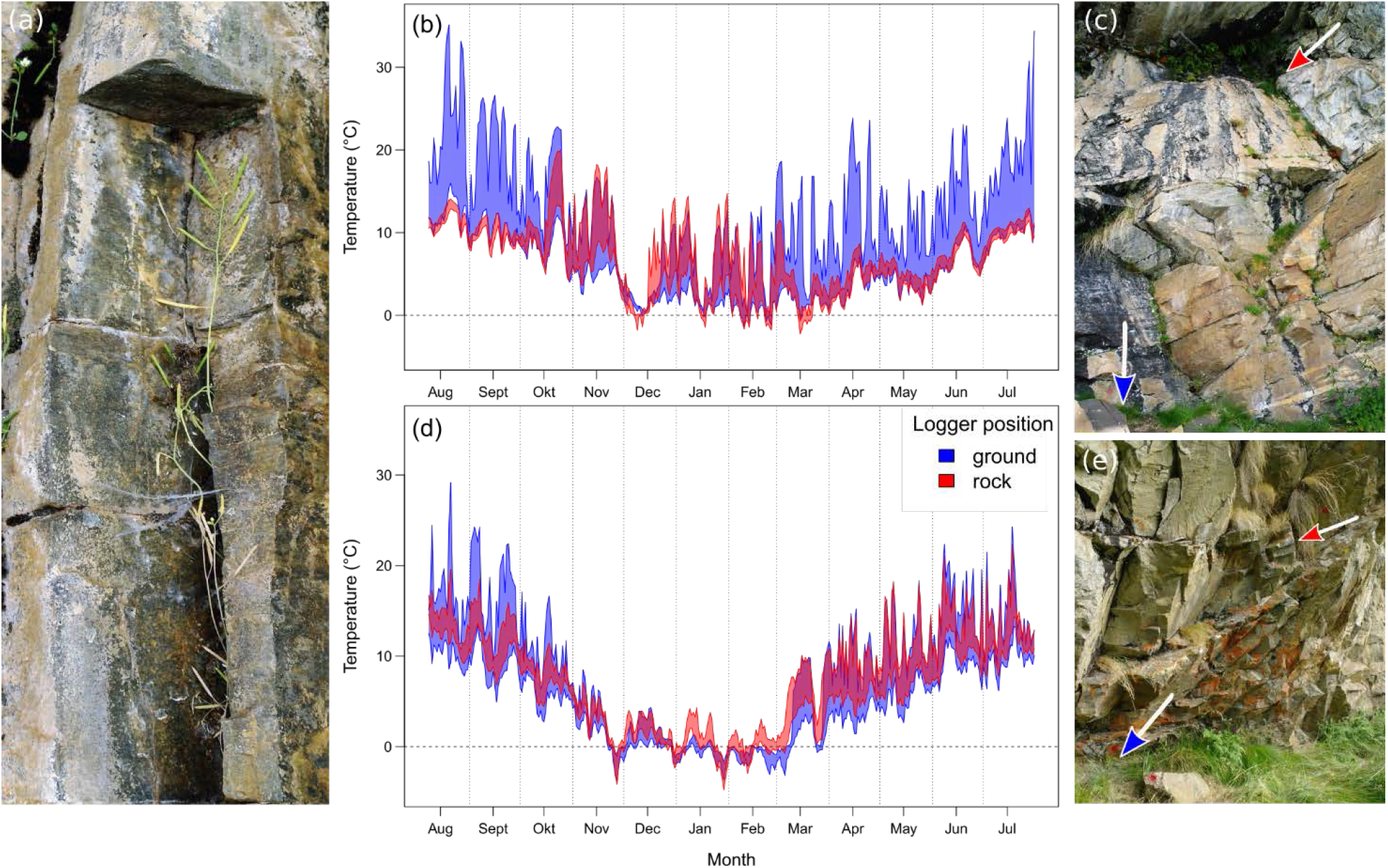
Top-soil temperatures across the year in two microhabitats of *A. thaliana* in the two highest populations. (a) *A. thaliana* plants growing in rock cracks at the site Cima Vioz. (b) Minimum and maximum top-soil temperatures of the two microhabitats rock and ground in Cima Vioz with the logger positions indicated by colored arrows in (c). (d) Minimum and maximum top-soil temperatures in Coronaccia with the logger positions indicated in (e).

## Discussion

### Frost-hardiness of Arabidopsis thaliana increases with altitude

In line with our initial prediction, frost-hardiness increased on average with the altitude of the collection site. This result is consistent with the well-documented difference in frost-hardiness between low-altitude and high-altitude species (Earnshaw *et al*., 1990; Taschler & Neuner, 2004). However, with respect to intra-specific variation in frost-hardiness the existing literature is more ambiguous. Sierra-Almeida et al. (2009) found higher frost resistance of populations from higher altitudes in four out of seven studied species from the high Andes. In the three remaining species frost-hardiness did not differ between high and low altitudes. Frost-hardiness also increased with higher altitude in the fern *Blechnum penna marina* from New Zealand (Bannister & Lee, 1989) and in *Solanum acaule* from Peru (Li *et al*., 1980). *Trifolium repens*, on the other hand, showed no differences in frost-hardiness between low- and high-altitude populations in Sweden (Junttila *et al*., 1990). These differences between studies and species are surprising given that frost damage entails a serious fitness cost (Agrawal *et al*., 2004). It suggests that altitude of provenance is not consistently a good proxy for frost-hardiness. Notably, in our study a log-linear curve greatly improved the model, suggesting that frost-hardiness changed rapidly with altitude at lower altitudes, but slowly at high altitudes. Indeed, frost-hardiness did not increase much above 1000 m despite strong variation among populations. This observation reflects our results in an earlier study with *A. thaliana* populations from the same region in South Tyrol, in which the differences in frost-hardiness among populations did not co-vary with the altitude of the five collection sites (Günther *et al*., 2016). In the present study we included more populations from lower altitudes, which improved the statistical power of the analysis of the relationship between frost-hardiness and altitude. Besides the variation among populations we also observed differences among years. On average these differences matched the differences in severe frost days at the common garden site on the Swabian Jura between the three experiments. However, beyond these differences also some individual populations varied strongly between the first two experiments, for example at Altenburg, Bozen-Train, Schnatz or Cima Vioz (Fig. 3). These differences may have several reasons. First, the experimental design differed among years, with some populations being represented by different accessions in different years (Table 1). However, two populations that were among the group showing the strongest variation (Bozen-Train and Schnatz) were always represented by the same accessions throughout the experiment. Therefore, we suggest that these changes among years may be rather attributable to differences in frost-acclimation. The range of temperatures that precede a frost event is very important for plant survival (Thomashow, 1999). Since other populations like Castelfeder, Vetzan and Finail showed nearly the same frost damage in the first two experiments (Fig. 3), our results indicate that populations vary not only in frost-hardiness, but also in their specific requirements for acclimation conditions. This has been observed previously for frost-hardiness in *Trifolium repens* (Junttila *et al*., 1990). Taken together, we observed high variation among populations and experiments that was partly associated with altitude. One reason for the high tendency of provenances to depart from the linear prediction for their altitude may be that frost-hardiness is more closely connected to the local microclimatic conditions than to the general climatic conditions at a specific altitude level.

### Probability of ground frost explains frost-hardiness better than altitude

We found that the frequency of ground frost estimated from top soil temperatures at collection sites, was superior to altitude as predictor of frost-hardiness differences between *A. thaliana* populations. In contrast to altitude, the frequency of frost days showed a linear relationship with frost-hardiness. This is in line with our second prediction and confirms that the microclimate rather than the altitude of a site accounts for the frost-hardiness of local populations. Mountains are characterized by a high degree of micro-topographic differences that influence the local microclimate (Briceño *et al*., 2014; Lembrechts *et al*., 2018). In particular, local frost conditions play an important role in microclimate adaptation. Wos and Willi (2018) showed that genotypic differences in frost-hardiness of *Arabidopsis lyrata* were linked to vegetation cover on a scale of a few meters in a sand dune landscape. In alpine environments, populations of *Aciphylla glacialis* from sites with early snow melt showed stronger frost-hardiness than populations from sites with later snow melt (Briceño *et al*., 2014). Also, frost damage of flower buds from three mountain wildflower species was associated with differences in snow accumulation, snowmelt pattern and cold air drainage on a scale of few meters (Inouye, 2008). Together these studies provide substantial evidence that microclimatic conditions independently of altitude affect plant growth and fitness. Furthermore, the mild microclimate of south exposed sites in alpine ecosystems supports plant growth and establishment of species whose core distribution range is at much lower altitude (Lembrechts *et al*., 2018). Our focal species, *A. thaliana* can be seen as a typical example for this phenomenon. As annual plant, it shows a life history that is strongly underrepresented in high alpine plant communities (Körner, 1995). In the present study *A. thaliana* populations occupied only SE to SW exposed slopes at high altitude (Table 2). In conclusion, microclimatic heterogeneity is strong in alpine environments and may allow species to occur at higher altitude than would be expected based on their ecological niche.

### Ground frost probability increasingly varies with higher altitude

We observed an increasing variance in top-soil temperature with increasing altitude of the site. Some high-altitude sites showed a similar number of frost days as low altitude sites, suggesting that at high altitude some *A. thaliana* populations exist in favorable microrefugia, which is consistent with the “law of the relative constancy of habitat” of (Walter & Walter, 1953). Such microrefugia are not uncommon in high mountains (Dobrowski, 2011b; Graae *et al*., 2012). According to Graae et al. (2012), the local temperatures in high mountain sites, when measured *in-situ*, are mostly higher than expected from interpolations across weather stations. The authors attributed this effect to inverted temperatures in winter, when cold air downdrafts hinder the accumulation of cool air at high-altitude sites. In line with this suggestion, all high-altitude sites in our study were situated on steep predominantly south-exposed slopes well above the cold air drainage that must be expected in the couloirs. Microclimatic conditions may also be influenced by differences in the effective heat capacity of soil and base rock. Accordingly, we found strong differences in temperature profiles between soil in rock-cracks and the soil below the rocks, which represent two microhabitats that were occupied by *A. thaliana* in the two highest sites. Specifically, the soil temperature in rock cracks showed reduced variation in summer, which can serve to buffer extreme heat peaks. However, also the most frost-hard population Finail was one of the high-altitude sites, which demonstrates that *A. thaliana* was not restricted to warm microrefugia at high altitudes. Besides habitat sorting according to the law of Walter and Walter (1953), also adaptation in frost-hardiness played a role in the successful survival of *A. thaliana* at high-altitude sites. In this population plants were found underneath larch trees which intercept snowfall reducing the snow depth beneath their crown. Snow layers are known to function as insulating layers that efficiently reduce frost damage (Inouye, 2008). This also matches the observation that frost-hardiness was associated with vegetation cover in *A. lyrata* (Wos & Willi, 2018). The potential reasons for microclimate heterogeneity are multifold. Together, our results suggest that at high altitude microclimate effects on top-soil temperatures may overcome average altitude effects. This was also suggested by Shreve (1924), who compared soil temperatures of north and south slopes at different altitudes. However, in contrast to Shreve (1924) who chose representative sites for each altitude, our sites mark actual populations of a species and we show that the microclimate had strong effects on local plant adaptation. This highlights two aspects that may be important for the survival of some plant species in the face of climate change. First, the microclimate heterogeneity of high alpine environments may preserve a high functional genetic diversity in small populations. Genomic results from our previous study suggested that high and low altitude populations of *A. thaliana* from South Tyrol split before the last glacial maximum, ca. 18.000 years ago (Günther *et al*., 2016). Second, the observed microclimate heterogeneity and its effects on the persistence of differently adapted populations at high altitude suggests, that using large scale climatic parameters to predict the fate of a species in a global warming scenario, as is common practice in climate-envelope modeling, may seriously underestimate the suitable habitat that is available in topographically complex regions.

## Aknowledgements

We thank the station chief of the Oberer Lindenhof agricultural station Helmut Bimek for excellent support with the common garden experiment the technical staff Oberer Lindenhof for help with the experimental setup. Anna Lampei Bucharova for help with the data logger collection in South Tyrolia. This work was supported by the F. W. Schnell Endowed Professorship of the Stifterverband to K. J. S.

## Author Contributions

C.L. and K.J.S conceived and designed the study; J.W., T.W., K.J.S and C.L. collected the seeds, C.L. conducted the experiments, performed the statistical analysis and wrote the first version of the manuscript, with J.W., T.W. and K.J.S contributing to revisions.

