## Supplemental Figure 1 for "Microclimate predicts frost-hardiness of alpine *Arabidopsis thaliana* populations better than altitude because the microclimate effect increases with altitude"

### Supporting information

Article title: **Increased microclimate effects at high altitude on the adaptation to local frost frequency in Southern Alpine Arabidopsis thaliana populations**

Authors: Christian Lampei[1,2], Jörg Wunder[3], Thomas Wilhalm[4], Karl J. Schmid[1]

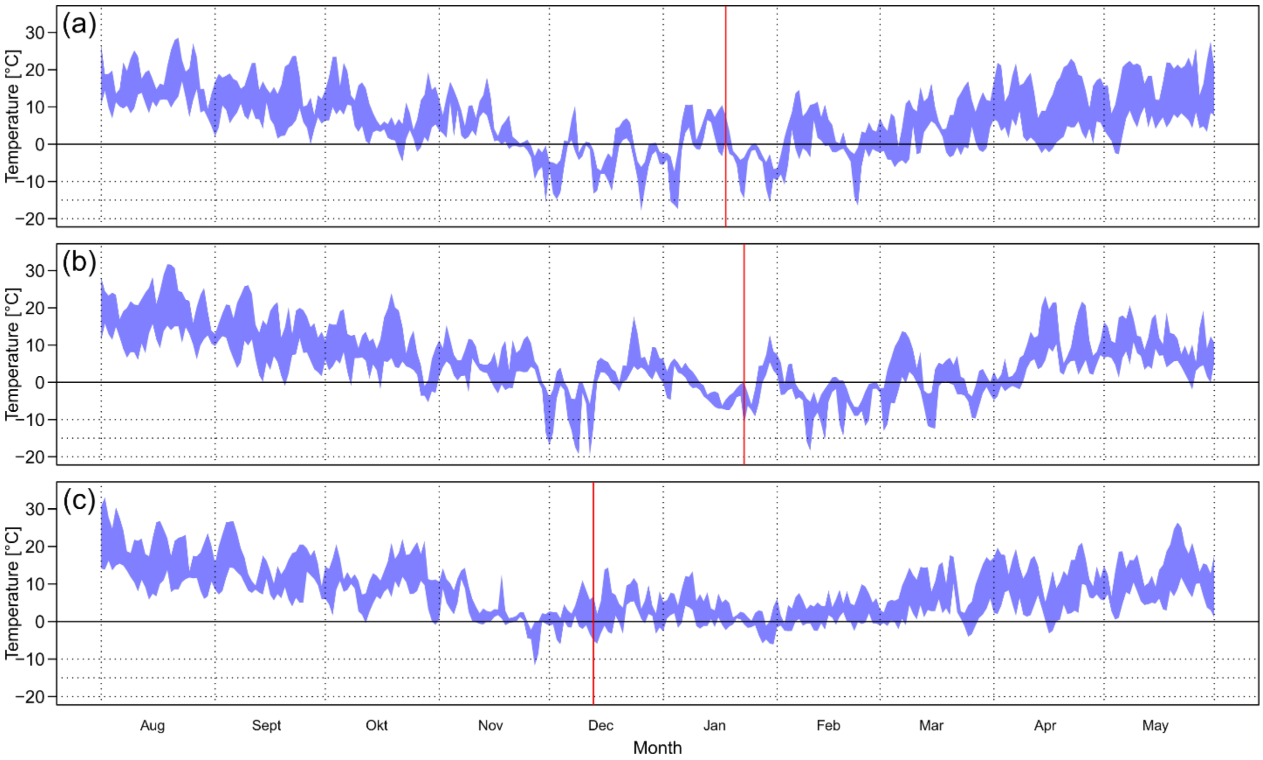


**Figure S1** Minimum and maximum temperature in the experimental site for the winter (a) 2010/2011, (b) 2012/2013 and (c) 2013/2014 obtained by the weather station of the experimental site. The red vertical line indicates the date when frost damage was evaluated. The number of days below 0 °C before the evaluation was (a) 55, (b) 52 and (c) 24. The number of days below -7 °C before the evaluation was (a) 18, (b) 14 and (c) 2. Due to technical problems, no weather data was available for February and March 2014. For this time the next neighboring weather station at similar altitude was used (St. Johann, 3 km distance at 749 m altitude).
